# Multiple forms of sensory reinstatement in category-selective cortex

**DOI:** 10.64898/2026.08.10.743957

**Authors:** Deepasri Prasad, Adam Steel, Caroline E Roberston

**Affiliations:** Dartmouth College, Hanover, New Hampshire 03755 USA; University of Illinois Urbana-Champaign, Champaign, Illinois 61820 USA

**Author notes:** **Author Contributions:** A.S. and C.E.R. designed research; A.S. and D.P. performed research; D.P. analyzed data; D.P. wrote manuscript; A.S. and C.E.R. edited manuscript.

**Keywords:** anterior shift, fMRI, visual perception, visual memory

## Abstract

Visual recall is classically thought to depend on reinstatement: areas engaged when encoding a visual input are similarly reactivated when remembering it. Here we investigated if reinstatement might be differently implemented across the diverse category-selective systems of visual cortex. Using fMRI in 25 participants, we assessed possible reinstatement organizations across scene-, face-, and body-selective cortex. We asked whether memory reactivates the same category-selective areas engaged during perception, whether it engages same or distinct vertices, and whether perceptual-mnemonic distinctions were topographically organized. All regions were selectively engaged during both perception and memory, though memory activity was weaker overall. At the vertex-level, most regions—including body-selective LOS, ITG, MTG; face-selective FFA1, FFA2; and scene-selective PPA—showed classic reinstatement, with memory enriched in the most perceptually selective vertices. In contrast, OFA and OPA showed separable perception-and memory-biased vertices. Critically, only scene-selective areas showed topographic distinction: in both PPA and OPA, mnemonic activity was located consistently anterior to perceptual activity, whereas no face-or body-selective areas showed such a distinction. Thus, while all category-selective areas are reactivated during memory, scene-selective cortex topographically separates memory from perception, suggesting different sensory reinstatement implementations across high-level visual cortex, possibly reflecting the distinct computational demands.

## Introduction

How the brain reconstructs our rich mnemonic representations in the absence of perceived sensory input is central to understanding memory (Goldberg et al., 2006; Pearson, 2019). Early work revealed that category-selective visual cortices, like place-selective parahippocampal place area (PPA) and face-selective fusiform face area (FFA), are not only active during the perception of their preferred visual category, but also when remembering that same visual category (O’Craven & Kanwisher, 2000). This idea of sensory reinstatement has been foundational to the study of visual memory, with extensive subsequent studies demonstrating area-level reactivation across a variety of visual categories, including areas in high-level visual cortex selective for processing scenes, faces, objects, and body parts (Hofstetter et al., 2012; Ishai et al., 2000; Ishizu et al., 2009; Johnson & Johnson, 2014; Khuvis et al., 2021; Polyn et al., 2005; Slotnick, 2004; Wadia et al., 2026). However, this focus on whether visual areas are reactivated during memory has left unclear how mnemonic activity is organized relative to perceptual activity within areas, and whether this organization might differ depending on the content being remembered (Favila et al., 2020, 2022; Breedlove et al., 2020; Steel et al., 2021; Y.Y. Chen, 2023).

Given the varying representational demands across high-level visual areas, it is not obvious that each system would organize mnemonic activity in the same way. Scenes in particular pose a distinctive challenge for perceptual-mnemonic integration: scenes are immersive, and scene understanding requires knowing not only the view of the scene that is perceivable, but also knowledge of the scene portions that are immediately out-of-view (Intraub, 2012). Comprehension of faces and bodies also depend on memory, but their visual analysis is more often organized around bounded agents, including information about identity, familiarity, expression, pose, and action. These differences motivate the possibility that perception–memory relationships may differ across scene-, face-, and body-selective cortex.

Recent work has revealed a few ways in which perception–memory relationships can vary at a finer scale than is captured by regional reactivation alone. One example is local intermixing, in which perception-and memory-biased responses coexist within the same cortical area, but differ across cortical layers. In V1, for instance, perceived color is decodable across cortical layers, whereas imagined color is preferentially decodable from deep layers (Bergmann et al., 2024). Similarly, single-neuron recordings within face-selective areas in both human (Quian Quiroga et al., 2023) and non-human primate (She et al., 2021) reveal heterogeneous familiarity effects, with individual neurons varying in their responsiveness to familiar and unfamiliar faces. A second possibility is topographic separation, in which mnemonic activity is systematically displaced from perceptual activity along the cortical surface. For example, visual recall of scenes elicits activity immediately anterior and adjacent to perceptual scene-selective areas, with partial overlap between perception and memory responses (Steel et al., 2021, 2023; Bainbridge et al., 2021; Srokova et al., 2022). By contrast, face-selective cortex does not appear to show the same topographic separation between perception and memory (Steel et al., 2021; Y. Y. Chen et al., 2024). Together, these findings suggest that perceptual-mnemonic transformations might assume distinct spatial organizations across fine-grained or regional scales within visual cortex.

Here, using individualized, within-subject fMRI, we test three possible organizations of mnemonic reactivation across independently localized scene-, face-, and body-selective regions spanning ventral and lateral occipitotemporal cortex (Fig. 1A). First, we examined the vertex-level correspondence between perception and memory. Classic reinstatement predicts that the most active vertices during perception should similarly be the most active during memory. An intermixed organization would predict that perception-and memory-biased vertices are separable but locally interspersed within the same category-selective area. Next, we assessed the spatial arrangement of perception and memory activity. A topographic distinction between the two tasks would predict that mnemonic activity is systematically displaced from perceptual activity along the cortical surface. It is important to note that vertex-level correspondence and spatial organization are independent: the same vertices may respond overall during both perception and memory activation in an area, even if the peak of the overall mnemonic response is nonetheless spatially displaced from the perceptual response. In brief, we find that category-selective regions are broadly reactivated during memory, but that the spatial organization of this reactivation differs across systems: only scene-selective areas demonstrate a topographic distinction between perception and memory.

**Figure 1.**
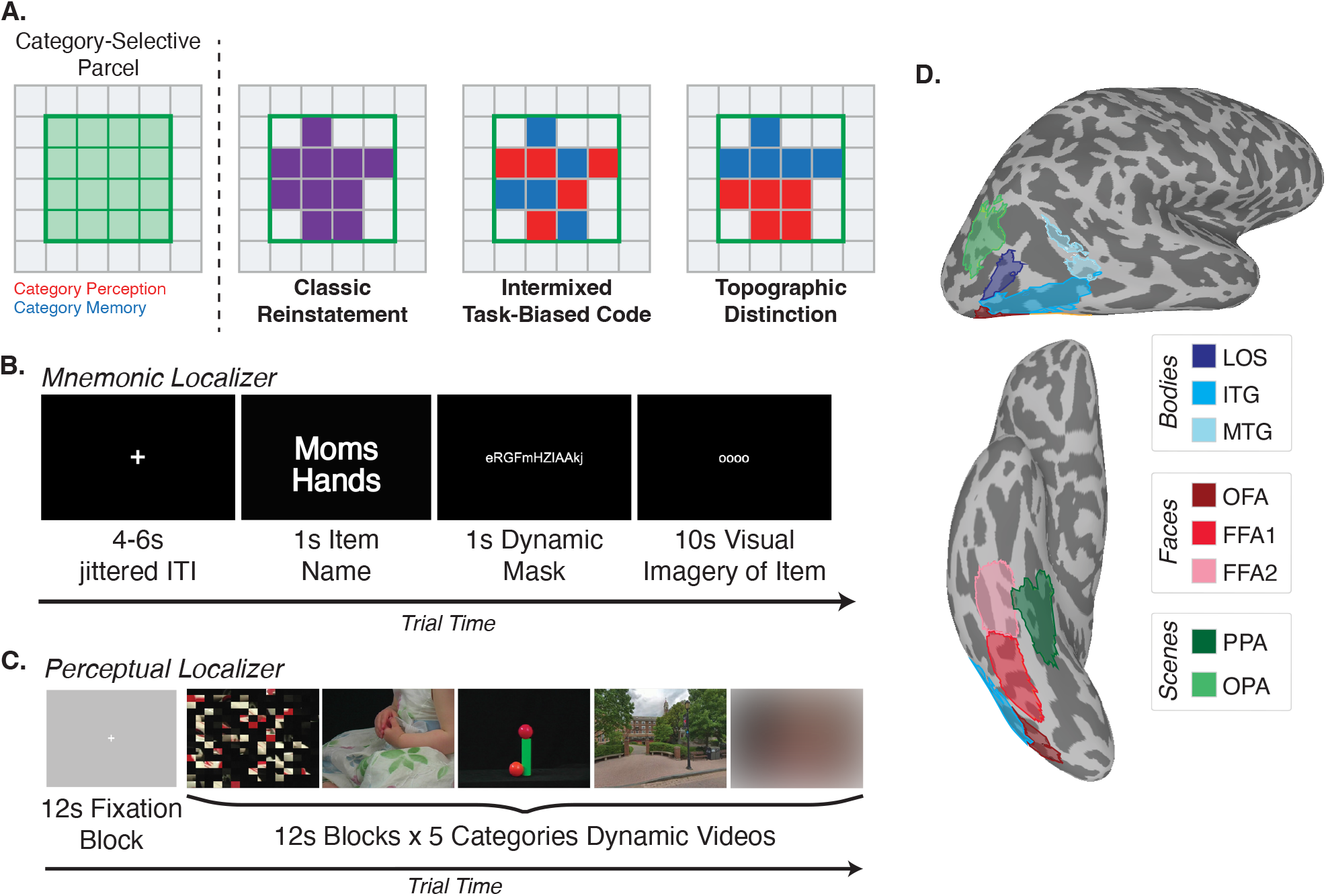
Candidate Implementations of Sensory Reinstatement and Definition of Category-Selective Areas. (A) Schematics depicting three tested organizations of mnemonic reactivation: *classic reinstatement*, in which memory reactivates the same perceptually selective vertices; an *intermixed code*, in which separable perception-and memory-biased responses coexist locally; a *topographic distinction*, in which memory responses are topographically distinct from perception. (B) Mnemonic localizer: participants viewed an item name from list of self-generated personally familiar faces, scenes, objects, and body parts (1 s), followed by a dynamic alphanumeric mask (1 s) and a visual imagery period (10 s) during which they vividly imagined the item. (C) Perceptual localizer: participants passively viewed 12s blocks of dynamic videos from five categories (scenes, faces, body parts, objects, scrambled). (D) Category-selective regions of interest were defined for each participant by constraining the t-statistics from participant-level category contrasts (category of interest > all other categories) with independent functional parcellations (Rosenke et al., 2021; X. Chen et al., 2023).

## Materials and Methods

### Participants

Twenty-eight adults were recruited for participation in the study. Three participants were excluded from analysis due to excessive motion artifacts, leaving twenty-five participants (17 females, age = 25.9 ± 4.2 STD years old) used in the analysis. Written informed consent was obtained from all participants. The study was conducted in accordance with relevant regulations, with a protocol approved by the Dartmouth College Committee for the Protection of Human Subjects (CPHS).

### MRI Acquisition

Data were collected on a Siemens Prisma 3T MRI scanner with a 32-channel head coil. Images were converted from DICOM to NIfTI files using dcm2niix, which by default applies slice time correction. For anatomical registration, a T1-weighted MPRAGE imaging sequence was acquired (TR = 2300 ms, TE = 2.32 ms, inversion time = 933 ms, flip angle = 8°, FOV = 240 × 240 mm, slices = 192, voxel size = 0.9 × 0.9 × 0.9 mm). Surface projections were generated from the T1-weighted images using FreeSurfer.

A multi-echo T2*-weighted sequence was used for all functional data (TR = 2000 ms, TEs = [14.60, 32.84, 51.08 ms], GRAPPA = 2, flip angle = 70°, FOV = 240 × 240 mm, Matrix size = 72 × 90, slices = 52, Multi-band factor = 2, voxel size = 2.7 × 2.7 × 2.7 mm). The slices were oriented parallel to the temporal lobe. The initial two frames were discarded by the scanner.

### Procedure-fMRI General

Participants underwent two fMRI localizer tasks: an mnemonic task to localize category-selective activity during visual recall (six runs, 48.8 min total; adapted from Steel et al., 2021) followed by a perceptual task to localize category-selective activity during visual perception (six runs, 41.0 min total; adapted from Pitcher et al., 2011).

### Procedure-Mnemonic Localizer

Participants provided a list of personally familiar places (e.g., their kitchen), personally familiar faces (e.g., their mother), personally familiar objects (e.g., their laptop), personally familiar body parts (e.g., mother’s hands), and known famous faces (e.g., Barack Obama) to be used as stimuli during the mnemonic localizer task (five items per category, twenty-five items total). Stimuli were generated prior to scanning, using specific guidelines for each visual category (See *Supplementary Materials*).

Outside of the scanner, participants completed a short practice that detailed how to perform the visual imagery during the task:

“*In this experiment you will be imagining people’s faces, places, bodies, and objects. On each trial, you will see either the name of a person, place, body, or object, followed by some scrambled letters and then 4 white circles. During the 4 white circles (10 seconds) your task is to imagine the item in as much detail as possible. Remember to keep your eyes open while imagining the item*.

### Objects

*For personally familiar objects, imagine that the object is turning around so you see all of its sides. Imagine the object without any context (i.e. no background). So, you will just be thinking about the object in front of you with nothing around it, just floating in space and turning around back and forth for 10 seconds (example on the right)*.

### Faces

*When imagining a famous person or personally familiar person, imagine their face in as much detail as possible and imagine their face is turning so you’re also able to see the side of their face. You can imagine them making different faces or emotions. It is important that you imagine their face visually*.

### Places

*When imagining a familiar place, imagine you are looking around in that place even though you are not physically turning your head. Picture the scene completely surrounding you and look around just as you do in real life. Do not walk around the place. Just imagine that you are in one spot looking all around you*.

### Bodies

*When you imagine familiar bodies, imagine only that specific body part. You can imagine that body part moving and turning so you can see all of its sides. It is crucial that you do not imagine the face of the body.”*

For each trial, participants were presented with an item name from their provided list for 1 s (Fig. 1B). After a 1 s dynamic alphanumeric mask, participants were instructed to visually imagine the item in as much detail as possible for 10 s. Each run included twenty-five trials (one trial per item, 4-6 s jittered ITI), with no more than two instances of a stimulus category appearing consecutively. The task consisted of six runs in total, and all participants completed all runs.

### Procedure-Perceptual Localizer

Participants viewed dynamic videos of five visual categories: scenes, faces, body parts, objects, and scrambled (Fig. 1C). Each category was presented in 12 s blocks, consisting of four 3 s-long videos, shown continuously. All category blocks were presented in a random order before showing a 12 s fixation rest block. Category blocks were then re-randomized and presented again, with each category repeated 5 times per run. Participants were instructed to watch the videos passively, with no task response solicited. Six runs in total were presented. Twenty-three participants successfully completed all six runs, with two participants successfully completing five runs.

### Preprocessing

Preprocessing was implemented using AFNI, based on the multi-echo preprocessing pipeline (afni_proc.py). Slice timing correction (3dTshift) and signal outliers were attenuated (3dDespike) separately on each echo. The second echo was used to estimate motion correction parameters (3dVolreg); these alignment parameters were applied to all echoes. Data was then denoised using multi-echo independent components analysis (ICA) (tedana.py). The optimal combination of the three echoes was calculated, and the echoes were combined to form a single, optimally weighted time series (T2smap.py). Principal component analysis (PCA) was applied to the data, and thermal noise was removed using the Kundu decision tree, which specifically targets and removes components that explain a small amount of variance and do not have a TE-dependent signal decay across echoes. Subsequently, ICA was performed to separate the time series into independent spatial components and their associated signals.

These components were classified as signal and noise based on known properties of the T2* signal decay of BOLD versus noise. The retained components were then recombined to construct the optimally combined, denoised time series. Following denoising, images were blurred with a 3 mm Gaussian kernel (3dBlurInMask).

### Defining Category-Selective Regions of Interest

To define category-selective perceptual activity, the perceptual localizer data was modeled by fitting gamma function of the trial duration with a square wave for each condition (scenes, faces, bodies, objects, and scrambled) using 3dDeconvolve. For category-selective memory activation, the mnemonic localizer was also modeled by the gamma function of the trial duration for trials of each condition (familiar faces, places, objects, body parts, and famous faces) using 3dDeconvolve. Famous faces trials were modeled but not included in the subsequent contrasts and analyses. For both tasks, estimated motion parameters were included as additional regressors of no-interest. Category-selective contrasts were obtained for both tasks by using a general linear test comparing the coefficients of the GLM during the category of interest contrasted against the coefficients during all other categories. These contrast maps were then transferred to the SUMA standard mesh (std.141) using @SUMA_Make_Spec_FS and @Suma_AlignToExperiment.

We then constrained each contrast map using probabilistic functional parcels from two published atlases (X. Chen et al., 2023; Rosenke et al., 2021) to define our category-selective regions of interest. Specifically, surface parcellations were taken from the visfAtlas (probability threshold=0.20; scenes: parahippocampal place area (PPA), occipital place area (OPA); bodies: lateral occipital sulcus (LOS), inferotemporal gyrus (ITG), middle temporal gyrus (MTG); faces: occipital face area (OFA)) (Rosenke et al., 2021) and from the Chen et al. (2023) atlas (probability threshold=0.10; faces: fusiform face area 1 (FFA1) & fusiform face area 2 (FFA2)) (Fig. 1D). Importantly, we chose to consider FFA1 and FFA2 separately, because of prior work suggesting that FFA2 may be more mnemonic in nature than FFA1 (Parvizi et al., 2012; Rangarajan et al., 2014; Schrouff et al., 2020). Furthermore, as face-selective areas are often not present in the left hemisphere, we chose to focus our analyses on the right hemisphere. For each participant and task, each surface parcel was intersected with the t-statistic map of the relevant category contrast (e.g., faces > all other categories) to define the ROI. Group t-statistic maps for perception and memory were calculated across participants for each category-selective contrast using 3dttest++.

## Results

Our overall aim was to determine how mnemonic activity is organized relative to perceptual activity within high-level category-selective visual cortex. Specifically, we asked whether perceptual and mnemonic activity is shared or distinct at the fine-grained vertex level, and whether they are topographically distinct at the area level. We first established that each category-selective region was reliably engaged and retained its expected category preference during both perception and memory. We then tested possible organizations of sensory reinstatement within each region: classic reinstatement, intermixed organization, and topographic distinction. We evaluated these hypotheses within each category-selective area.

### Independent parcels capture category-selective activity during perception and memory tasks

Participants performed two tasks: 1) a perceptual localizer, in which participants passively watched videos of dynamic faces, places, body parts, and objects, and 2) a mnemonic localizer, in which participants vividly imagined personally familiar items from the same visual categories (see Materials and Methods). We constrained the resulting participant-level activation maps using functional parcels from two independent, perception-defined probabilistic atlases (Fig. 1D; X. Chen et al., 2023; Rosenke et al., 2021). Before using these parcels to test the vertex-level organization of reinstatement, we validated that they captured the expected category-selective activity in both tasks.

At the group level, whole-brain analysis revealed that each parcel captured significant category-selective responses during both perception and memory, with the peak of activation falling within the relevant category-selective area (Fig. 2A). Thus, although the independent probabilistic parcels were defined using perceptual activity, they were suitable spatial constraints for evaluating activity in both our perception and memory tasks. For each participant and task, we defined an individualized ROI as the vertices comprising the intersection of each parcel with the relevant category contrast (e.g., FFA1: faces > all other categories). We next assessed whether these parcels also captured significant category-selective activity during both tasks at the individual-participant level. For each ROI, we extracted the vertices constituting the top 10% of t-statistic values for the relevant category contrast. We calculated the mean of the t-statistic values across these vertices, and tested whether it exceeded a two-tailed significance threshold corresponding to p=0.05 (t-statistic=1.962) (Fig. 2B). During perception, all individualized ROIs exceeded this threshold (all t(24)≥6.049, all p<0.001; *Table S1*). During memory, responses were significantly weaker than perception in every ROI (Fig. 2B; all paired t(24)≥5.205, all p<0.001; *Table S1*). Nevertheless, almost all ROIs also showed mnemonic activity significantly above threshold as well (all t(24)≥1.862, all p ≤ 0.040; *Table S1*), with the exception of LOS and MTG, which approached, but did not reach, the threshold (LOS: p = 0.051; MTG: p = 0.058).

**Figure 2.**
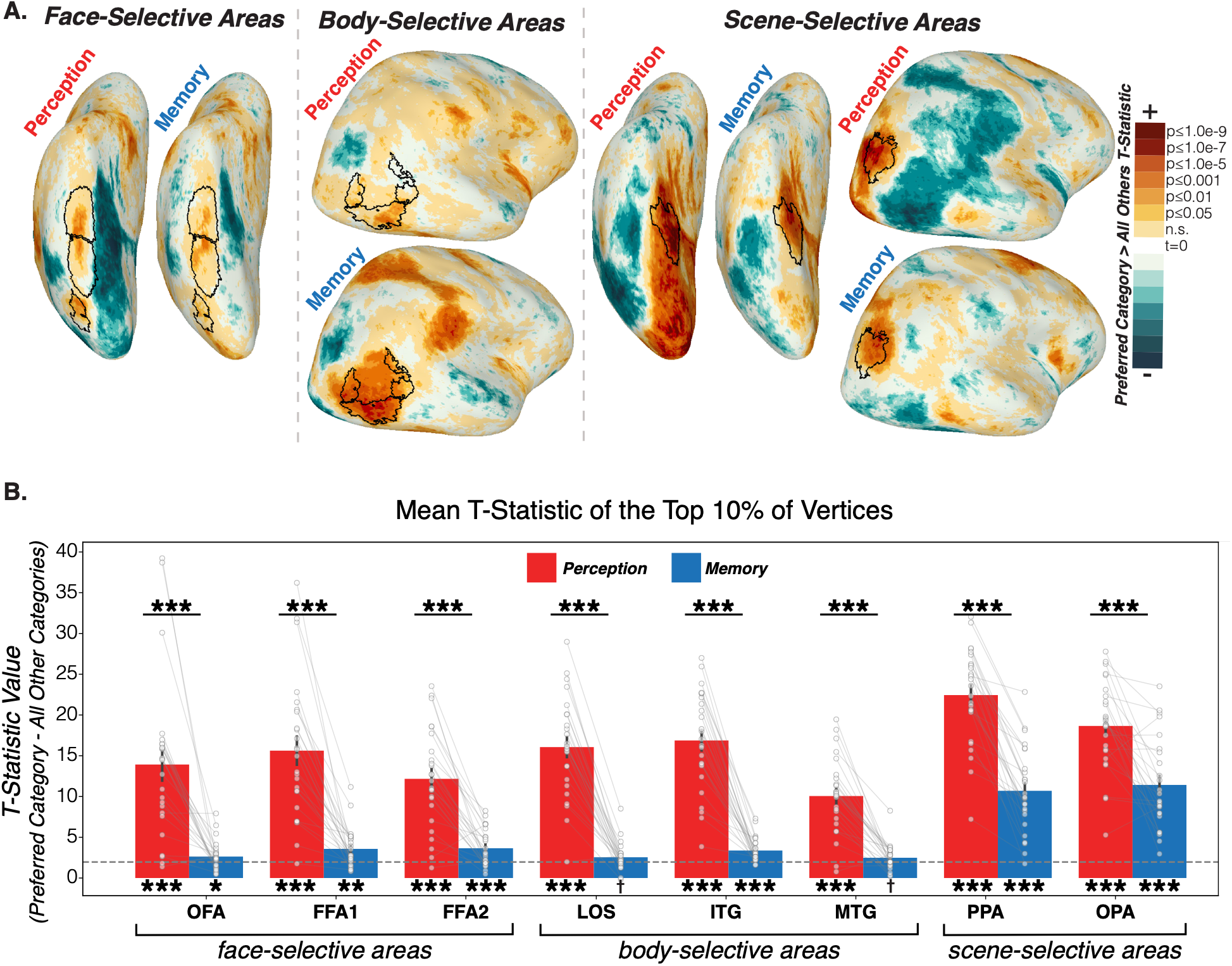
Category-selective area activity during both perception and memory. (A) Group-level t-statistic maps for each category contrast during perception and memory, showing peak activation contained within ROI boundaries for all categories. (B) Mean t-statistic of the top 10% most selective vertices within each ROI for perception and memory, tested against a threshold of p=0.05 (t=1.962; dashed line). All ROIs were significantly above threshold during perception, and almost all ROIs (with body-selective LOS and MTG approaching significance) were above threshold during memory. Activity was significantly lower during memory than perception across all ROIs. Error bars denote SEM; †p≤0.06, *p≤0.05, **p≤0.01, ***p≤0.001.

To visualize the responses underlying these category-selectivity omnibus contrasts (i.e., preferred category > all other categories), we then plotted the uncontrasted beta estimates for each category within each individualized ROI for each task (Fig. 3). The resulting profiles generally showed the expected category preference during both perception and memory, where all ROIs were significantly greater in their response for their preferred category compared to all other categories (Fig. 3A,B; one-tailed paired t-tests: all t(24) ≥ 2.236, all p ≤ 0.017; *Table S2*). One notable exception was ITG, which showed similar mnemonic responses to bodies and objects, consistent with its spatial overlap with lateral object-selective cortex (Grill-Spector et al., 2001; Weiner & Grill-Spector, 2011). The overall category-response profiles were also qualitatively similar across a range of vertex-selection thresholds (Fig. 3C). We additionally observed that the order of category preference remained the same across both tasks in most ROIs (Fig. 3A,B), except for OFA and MTG, which could suggest that ROI category preference order is a relatively stable property, regardless of the task. To validate that the vertex selection process was not biased towards the preferred category, we additionally performed a split half analysis, where the vertices to select were determined from half of the data, and category-selectivity was assessed on held out data. Qualitatively, this resulted in a similar trend of selectivity, although mnemonic selectivity was less specific for body-and face-selective areas (Supplementary Fig. S1).

**Figure 3.**
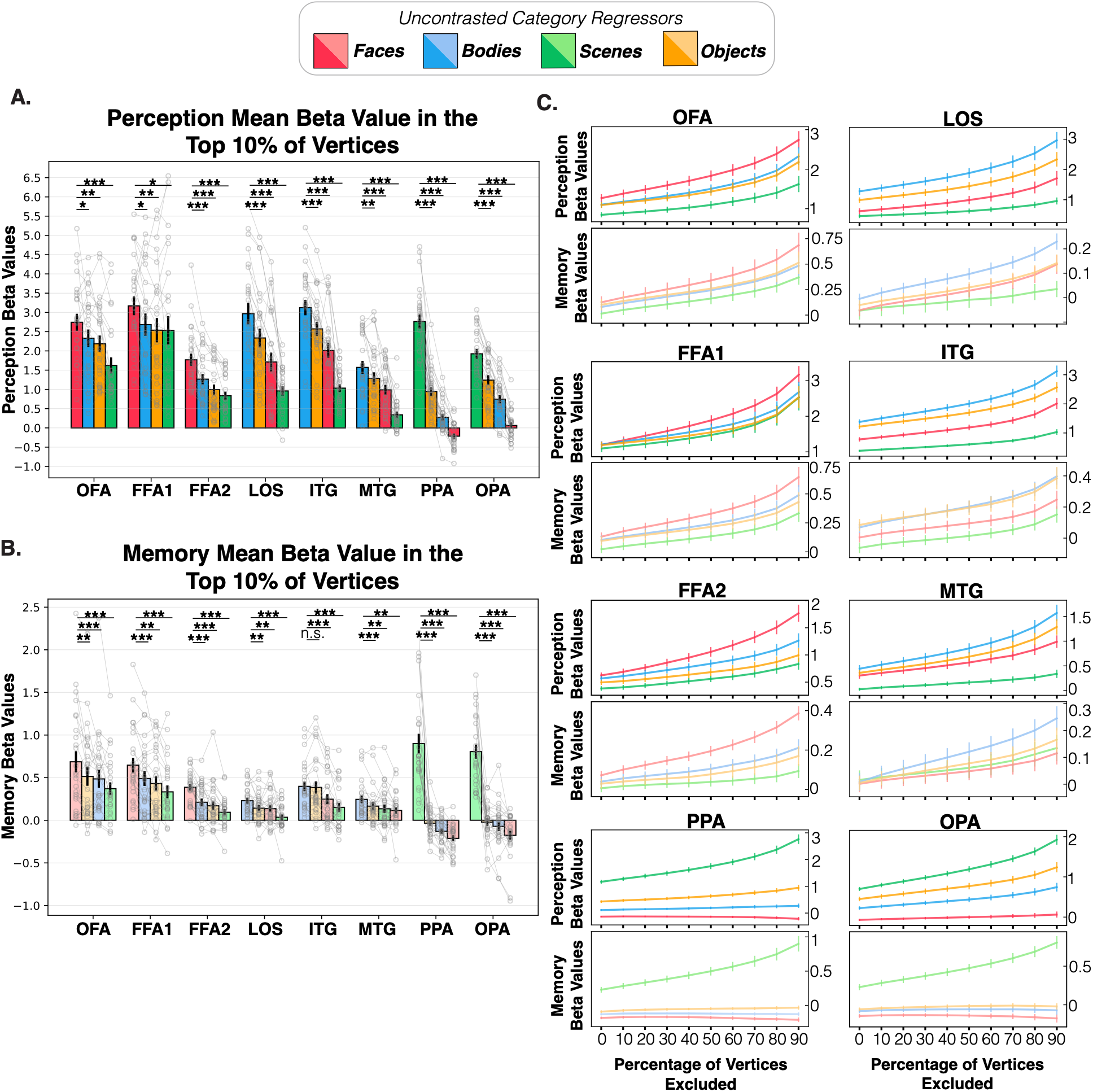
Category selectivity is maintained during both perception and memory. (A) Mean beta estimates to all four categories in the top 10% most responsive vertices to the preferred category for each ROI during perception, confirming that every ROI responded most strongly to its preferred category. (B) The same analysis during memory, showing preserved category selectivity in almost all ROIs (C) Selectivity for the preferred category across decreasing vertex inclusion percentages. Error bars denote SEM.

Together, these analyses provide coarse, area-level evidence for classic reinstatement: independently defined category-selective parcels retained their expected category preferences during memory, although mnemonic selectivity was weaker and less robust in some regions.

### Perceptual selectivity predicts mnemonic responses at the vertex level in most category-selective areas

We next asked whether this area level reinstatement extended to the fine-grained vertex level, or whether perceptual and mnemonic responses were separable and distinct within these areas. Within a classic reinstatement organization, the most selective vertices during perception should similarly be among the most active during memory. By contrast, within an intermixed organization, mnemonic activity need not be concentrated in the vertices that are most selective during perception, and perception-and memory-biased vertices may coexist within the same category-selective area.

To test these predictions, we stratified the vertices in each category-selective area based on their response during perception and compared the responses of those same vertices during memory. Under classic reinstatement, the mnemonic activity of the most perceptually selective vertices should be enriched compared to that of the least perceptually selective vertices, as the most and least selective vertices would correspond across the two tasks. An intermixed organization, however, predicts no such correspondence and therefore, would anticipate no systemic increase in the mnemonic response of the most perceptually selective vertices compared to the least. To investigate, we first ranked vertices within each ROI by their perceptual category-selectivity contrast and extracted the z-scored mnemonic response – that is, the category-selective contrast during memory – from the 20% most and least perceptually selective vertices. To examine the organization of jointly category-selective activity, we restricted the analysis to vertices with positive category-selectivity contrasts in both tasks.

Participants with fewer than 25 remaining vertices in the ROI were excluded. We then calculated the difference between the z-scored mnemonic response of the most and least perceptually selective vertices, where positive difference indicates that mnemonic responses were enriched in the most perceptually selective vertices.

Consistent with the classic reinstatement hypothesis, in most ROIs we found enriched mnemonic response in the top perception selective vertices, as shown by a significantly greater than zero difference (Fig. 4A). This included face-selective FFA1 and FFA2; body-selective LOS, ITG, and MTG; and scene-selective PPA (one-tailed t-tests: all t≥1.847, all p≤0.039; *Table S3*).

**Figure 4.**
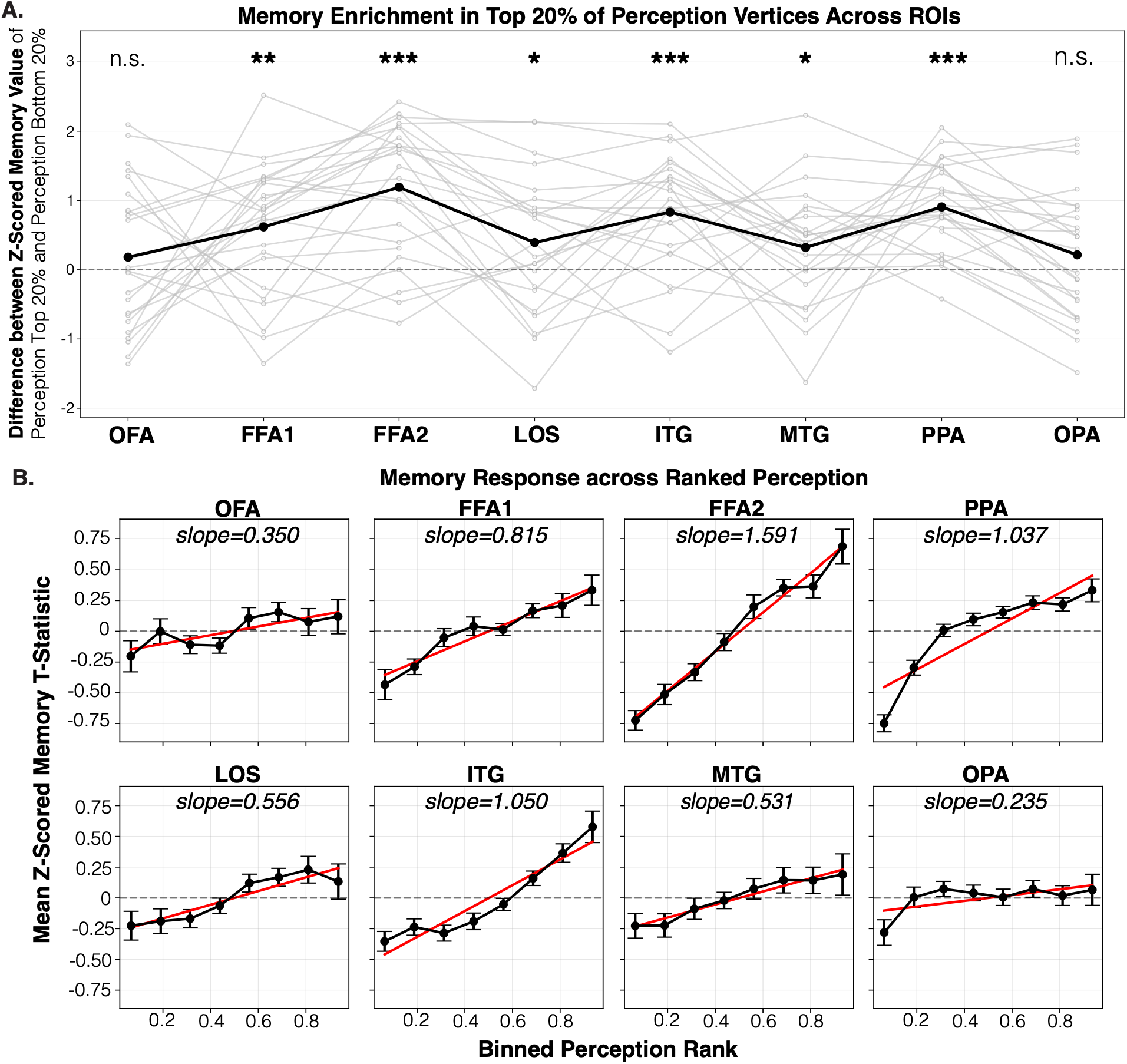
Vertex-level reinstatement in most, but not all, category-selective areas. (A) Difference in z-scored mnemonic activation between the 20% most and 20% least perceptually selective vertices within each ROI. Positive values indicate that memory activation is concentrated in the most perceptually selective vertices, as predicted by classic reinstatement. Enrichment was significant for FFA1, FFA2, LOS, ITG, MTG, and PPA, but not for OFA or OPA. (B) Mean memory activation across eight bins of perceptually selective vertices and the corresponding slope. Most ROIs showed a significantly positive slope, whereas OFA and OPA did not (OFA approaching significance). Error bars/shading denote SEM; *p≤0.05, **p≤0.01, ***p≤0.001

OFA and OPA were the only ROIs that did not show significant enrichment (OFA: t(23)=0.848, p=0.203; OPA: t(24)=1.171, p=0.127; *Table S3*), providing no significant evidence that their most perceptually selective vertices were preferentially responsive during memory. Thus, most category-selective regions showed vertex-level correspondence between perceptual selectivity and mnemonic responsivity, whereas OFA and OPA did not show significant enrichment.

Next, we examined whether this relationship was graded across the full range of vertices within the ROI, or whether it only applied to the topmost perceptually selective vertices. A positive slope would indicate that mnemonic responsivity systematically increased as perceptual selectivity increased, as predicted by the classic reinstatement model. In contrast, a flat slope would be more consistent with an intermixed model, in which mnemonic response was present within the ROI but not systematically aligned with perceptual selectivity. To test this, for each participant and ROI, we binned vertices by perceptual selectivity across eight bins, calculated the mean z-scored memory response for each set of binned vertices, and estimated the slope of memory response across these bins (Fig. 4B). Participants with less than 10 vertices in each bin for an ROI were excluded from the analysis (all ROIs ≥ 19 participants; Table. S4).

This analysis revealed a similar trend to the enrichment analysis: most ROIs showed a significantly positive memory slope, indicating a graded relationship between perceptual selectivity and mnemonic response across vertices (one-tailed t-tests: all t ≥ 2.08, all p ≤ 0.026; *Table S4*). OFA and OPA did not, although OFA approached significance (OFA: t(18) = 1.716, p = 0.052; OPA: t(24) = 1.025, p = 0.158; *Table S4*).

Together, these analyses provide evidence for vertex-level reinstatement in most category-selective areas: vertices showing stronger perceptual selectivity also tended to show stronger mnemonic responses. OFA and OPA were the exceptions, showing no evidence of preferentially enriched mnemonic activity in the most perceptually selective vertices, indicating that ROI-level reactivation in these regions was not organized according to classic vertex-level reinstatement.

### Mnemonic responses are anteriorly displaced relative to perceptual responses only in scene-selective cortex

Our results examining vertex-level correspondence between perception and memory demonstrated that most category-selective areas implement classic reinstatement at the vertex-level. We next investigated whether perceptual and mnemonic responses also differed in their large-scale topographic organization. This question is distinct from vertex-level correspondence: perceptual and mnemonic responses may substantially overlap within an area while nonetheless exhibiting a consistent displacement between their peak locations.

To test this, we compared the topographical coordinates of peak perceptual and mnemonic activation within each ROI, excluding any participant with peak activation below a threshold of p=0.05 (t=1.962). For each participant and ROI, we calculated the polar vector from the perceptual peak to the mnemonic peak on the cortical surface. We used the vector length to quantify the degree of peak separation and Rayleigh tests to assess whether vector directions were consistently oriented across participants. If the memory peaks are consistently oriented in a particular direction from their corresponding perceptual peaks, it would suggest a topographic distinction between the two processes.

Scenes uniquely showed a topographic separation between mnemonic and perceptual activity. For scene-selective PPA and OPA, the distribution of directions between perceptual and mnemonic peaks was significantly nonuniform across participants, with mnemonic peaks consistently located anterior to perceptual peaks (Fig. 5A; Rayleigh tests: PPA: Z=17.779, *p* < 0.001; OPA: Z=18.334, *p* < 0.001). The mean peak-to-peak distance was 15.084±1.336 SEM mm in PPA and 7.107±0.992 SEM mm in OPA. In PPA, this anterior displacement was present despite its significant vertex-wise reactivation: as visible in the group maps (Fig. 2A), perceptual activity extended farther posteriorly, and mnemonic activity extended farther anteriorly, whereas the strongest perceptual and mnemonic responses overlapped in anterior PPA. In OPA, the displacement provides a potential topographic explanation for the absence of significant vertex-level correspondence.

**Figure 5.**
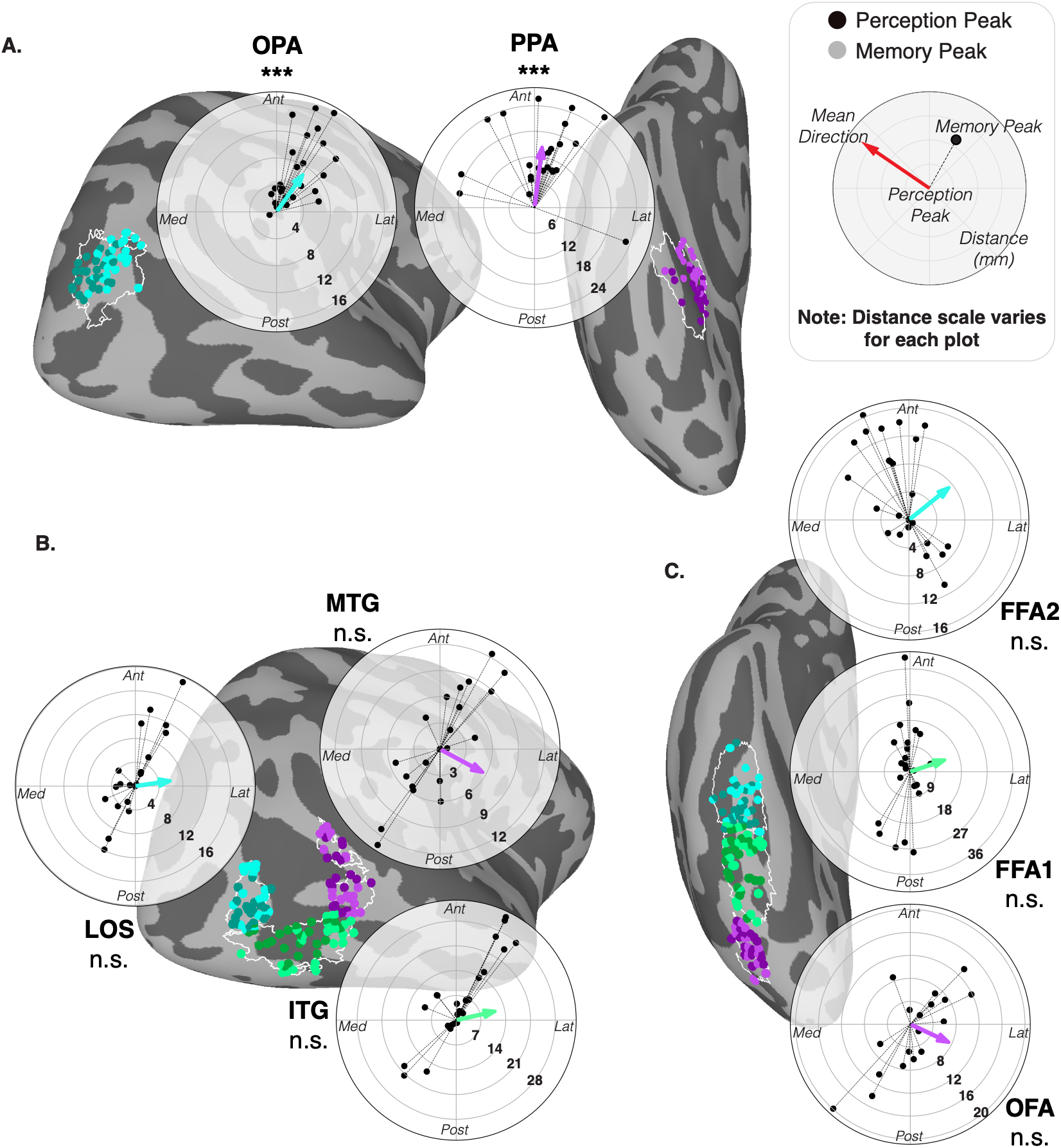
Only scene-selective areas show a topographic distinction between perception and memory. For each participant and ROI, we calculated a vector on the cortical surface from the peak perceptual response to the peak mnemonic response. Polar plots show individual-participant vectors (black points) and the mean vector (colored arrow); radial distance indicates peak-to-peak distance in millimeters. Perception peaks (dark colors) and memory peaks (light colors) for each participant plotted on surface, with each parcel boundary outlined (A) Scene-selective PPA and OPA showed significant topographic shifts of memory relative to perception, in the anterior direction. (B) Body-selective areas (LOS, ITG, MTG) showed no significant topographic distinction between perception and memory. (C) Face-selective areas (OFA, FFA1, FFA2) also showed no significant topographic distinction between perception and memory.

Importantly, we did not observe a significant topographic distinction between perception and memory in any other category-selective area. Body-selective areas did not show a significant topographic distinction between mnemonic and perceptual activity (Fig. 5B; Rayleigh tests: LOS:, Z=1.27, *p*=0.285; ITG:, Z=2.59, *p*=0.074; MTG:, Z=0.056, *p*=0.946). The mean peak-to-peak displacement was 6.230±1.119 SEM mm in LOS, 11.848±2.012 SEM mm in ITG, and 5.860±0.796 SEM mm in MTG. Such a topographic distinction was also not found within any of the face-selective areas examined, including OFA (Fig. 5C; Rayleigh test: OFA: Z=0.879, *p*=0.42; FFA1: Z=0.704, *p*=0.499; FFA2: Z=1.10, *p*=0.338). The mean peak-to-peak displacement was 7.777±1.068 SEM mm in OFA, 13.497±2.134 SEM mm in FFA1, and 7.736±0.996 SEM mm in FFA2.

Taken together, these results show that memory and perceptual activity are topographically distinct in scene-selective PPA and OPA, replicating the anterior shift for memory and perception observed in previous work (Steel et al., 2021, 2023; Bainbridge et al., 2021; Srokova et al., 2022). No consistent displacement was detected in the examined face-or body-selective areas, suggesting that topographic separation may be a distinctive organizing principle of scene reinstatement.

## Discussion

The classic sensory reinstatement account of visual recall predicts that the category-specific areas engaged when encoding a stimulus are reactivated during memory. Here we show that this principle broadly holds across scene-, face-, and body-selective cortex. Within a region, most areas showed vertex-level evidence that mnemonic responses corresponded to perceptually selective vertices. Critically, however, this reinstatement had a different spatial organization in scene-selective cortex. Both PPA and OPA showed a systematic anterior shift of mnemonic activity relative to perceptual activity, whereas face-and body-selective areas showed no comparable topographic separation. Thus, perceptual–mnemonic correspondence was common across category-selective cortex, but only scene-selective areas showed a consistent spatial transformation between perception and memory.

Our findings deepen traditional accounts of sensory reinstatement by showing that vertex-level correspondence is widespread across category-selective systems, while its broader topographic organization differs across systems. Classic reinstatement models predict that memory reactivates the same neural populations engaged during perception. However, most prior reports examine reactivation at the regional level (O’Craven & Kanwisher, 2000; Polyn et al., 2005), and studies testing finer-grained reactivation use only one or two visual categories (Wadia et al., 2026). By comparing multiple category-selective systems within vertex-level recruitment, we show that reinstatement is widespread but not uniform in its spatial organization. At the same time, the present analyses do not capture all possible forms of perception–memory transformation. For instance, it remains possible for transformations to involve intermixing of perception-and memory-biased substrates at a more granular level than can be resolved with this fMRI paradigm, such as at the level of cortical layers or individual neurons (Bergmann et al., 2024). Furthermore, decoding analyses would be sensitive to additional kinds of transformations between memory and perception, like those that occur along stimulus feature dimensions(Favila et al., 2020, 2022). Here, the perception and mnemonic localizers used different stimuli: unfamiliar dynamic images during perception and personally familiar stimuli during imagery. This design limited our ability to decode how the same stimulus transforms between perception and memory. Future studies should continue to examine reinstatement using matched perception and mnemonic stimuli across multiple visual categories, allowing stimulus-specific transformations to be compared directly across scene-, face-, and body-selective cortex.

The anterior displacement of mnemonic relative to perceptual activity in scene-selective cortex we observed replicates previous work showing that scene recall recruits regions anterior and adjacent to perceptual scene areas (Steel et al., 2021, 2023; Srokova et al., 2022). Indeed, previous work has also shown that peak mnemonic activation can even fall outside of and anterior to traditional scene perception areas, such as the ones used in the current study (Steel et al., 2021). What computational purpose might anteriorly shifted memory activation serve?

One possibility is that it reflects a broader posterior–anterior gradient, where posterior areas represent more perceptual, concrete information and anterior areas more conceptual, abstract information (Favila et al., 2020; Popham et al., 2021; Rugg & Thompson-Schill, 2013). Under this view, the spatial transformation between perceptual and memory representations would reflect a general principle by which memory representations become abstracted away from perceptual representations. However, if this were the explanation, similar anterior shifts might be expected within other category-selective systems. The absence of comparable topographic distinctions in face-or body-selective cortices suggests that a domain-general posterior– anterior abstraction gradient is unlikely to fully explain the scene-specific effect.

An alternative possibility is that this topographic shift reflects a distinctive computational demand of scene representation. Unlike other visual categories, scenes are immersive spatial environments: while a face or a body part may be fully captured by a single field of view, scenes by their nature contain not just what is immediately perceivable, but what is also currently out of view. Therefore, accurate scene representation may require close coordination between representations of the viewed portion of the scene and remembered out-of-view portions, while still maintaining distinction between perceptual and mnemonic representations (Steel et al., 2021, 2024). Consistent with this view, prior work has shown that memory activation for scenes extend anterior beyond the bounds of traditional scene perception regions, forming a “place memory network” (Steel et al., 2021, 2025). Activity in these anterior place-memory areas – but not their posterior counterparts – increases when more visuo-spatial context is known for a scene (Steel et al., 2023). These place memory areas also represent associations from panoramic viewing of scenes (Han & Epstein, 2026) and predictions of upcoming scene views (Mynick et al., 2026), consistent with a role in linking the current view to remembered spatial context. The adjacent but topographically distinct organization of scene perception and memory may therefore support the integration of the current view with remembered and anticipated spatial context.

The connectivity profile of scene-selective cortex also lends evidence to this interpretation. Posterior scene perception regions are more strongly coupled to early visual cortex, consistent with their role in representing the currently visible input. By contrast, anterior place memory areas appear more closely coupled to medial temporal and default-network regions, including the hippocampus, and form distinct functionally coupled networks across the cortical surface (Baldassano et al., 2016; Steel et al., 2021, Silson et al., 2016). These anterior areas may therefore occupy a transitional position between visual scene processing and broader memory systems. From this perspective, the anterior shift during scene recall reflects recruitment of cortical territory positioned to integrate the current view with prior experience and information beyond the field of view.

Other differences between the stimulus categories could also contribute. For example, personally familiar scenes may evoke broader contextual information, including geographic position of that location; multisensory information from that environment, like the sounds or the temperature; or even episodic events that have occurred at that scene. However, anterior mnemonic shifts have also been observed for recently studied, unfamiliar scenes lacking this broader personal context, indicating that personal familiarity is not necessary for the effect (Bainbridge et al., 2021; Steel et al., 2021). Scenes also typically extend farther into peripheral vision than faces or body parts (Park et al., 2024). Although images of scenes that do not extend into the periphery can recruit anterior place-memory areas (Bainbridge et al., 2021; Srokova et al., 2022) and peripherally-presented face and object stimuli induce expected activity in corresponding category-selective areas (Park et al., 2024), the contribution of eccentricity remains unresolved. Future studies could directly compare memory for peripherally and foveally presented stimuli across categories to determine whether peripheral engagement contributes to the distinctive topographic organization of scene recall.

Our findings in face-selective areas additionally replicate the findings in Steel et al., (2021) and Y. Y. Chen et al., (2024). Their work found that memory reactivation in OFA, FFA1, FFA2 (Y. Y. Chen et al., 2024), and combined FFA (Steel et al., 2021) overlaps with perceptual activation in those areas, with no topographic shift between perception and memory – just as we observed. Interestingly, however, while OFA did not show a topographic separation in the current study, it also did not show strong vertex-level reinstatement. Instead, mnemonic responsivity was not preferentially enriched in the most perceptually selective OFA vertices, suggesting the possibility of more separable perception-and memory-biased responses within this region. However, compared to the other regions, OFA had relatively small number of vertices, and overall did not show as much activity during memory, potentially limiting our ability to detect vertex-level reinstatement. More broadly, current evidence for memory reinstatement with faces has focused primarily on the more posterior face-selective regions. To our knowledge, no study to date has investigated topographic shifts in more anterior face-selective areas, including face patches in the anterior temporal lobe, which nonhuman primate and human research have implicated in memory (Collins & Olson, 2014; Deen et al., 2024).

Future studies should examine how reinstatement is implemented across the full posterior– anterior face-processing hierarchy, and whether tuning for perceived versus remembered faces changes along this axis.

We did not collect vividness ratings for individual trials in this study, leaving open the possibility that some differences we see across visual categories may be due to difficulty in imagining that category or low vividness of their imagery. However, all ROIs in our study showed significant category-selectivity during both perception and memory, mitigating that concern. Additionally, other work studying the condition of aphantasia, the lack of voluntary visual imagery, suggests that there is no difference in the degree of anterior shift for scenes memory between individuals who can perform visual imagery versus those who cannot (Megla et al., 2024). As such, it is unlikely that our findings are being driven by differences in the vividness of visual recall.

In summary, expected category preferences were broadly reinstated during visual memory, and most regions showed vertex-level correspondence between perceptual and mnemonic selectivity. Uniquely, however, scene-selective PPA and OPA showed a systematic anterior shift of mnemonic relative to perceptual activity. These findings suggest that sensory reinstatement may be implemented differently across high-level visual systems, potentially reflecting the specialized computational needs of each visual category.

## Acknowledgements

This work was supported by NSF Career Award 2144700 (C.E.R.) and NSF GRFP Award 2236868 (D. P.). We are grateful to all participants in our study.

