## Supplemental Materials for "Multiple forms of sensory reinstatement in category-selective cortex"

**Supplementary Materials**

*Mnemonic Stimuli Generation Instructions*

Participants were given the following instructions to generate the stimuli used during the mnemonic localizer task:

*"During the scan, you will be asked to visualize people, places, objects, bodies, and everyday objects that are personally familiar to you. So, we need you to provide these lists for us.*

*People: For people, you will be choosing 5 famous people you have not personally met before (e.g., Ariana Grande) and 5 personally familiar people that you know personally that you can visualize in great detail. You do not need to be in contact with these people now—just as long as you knew them personally and remember what they look like. So, you could choose your childhood friend even if you are no longer in touch with this person.*

*Places: Please list 5 personally familiar places that you have been to and can imagine in great detail. You should choose places that are personally relevant to you, so you should avoid choosing places that you have only been to one time, and you should not choose famous places where you have never been. You can choose places that span your whole life, so you could do your current kitchen as well as the kitchen from your childhood home.*

*Objects: For objects, choose 5 personally familiar objects that are about the size of a toaster (e.g., your favorite mug, a coffee maker). These objects should be personal items that you know very well and interact with fairly regularly (e.g., your personal hair brush or your soccer ball).*

*Bodies: For bodies, you will choose 5 familiar parts of bodies that are specific to someone you know (e.g., the hands or feet of your parents). The parts of the body you choose do not have to belong to the personally familiar people you chose. Please do not include explicit body parts. Some examples of body parts specific to a person could be hands, feet, legs, arms, whole bodies etc., but not faces."*

**Title:** Multiple forms of sensory reinstatement in category-selective cortex

*Supplementary Tables & Figures*

**Table S1.** Top 10% of vertices in perception and memory: One-tailed T-test against threshold and paired t-test

| *ROI* | *Perception Mean t-statistic value ± SEM* | *Memory Mean t-statistic value ± SEM* | *DF* | *Perception: one-tailed t-test against threshold of p=0.05 (t=1.962)* | | *Memory: one-tailed t-test against threshold of p=0.05 (t=1.962)* | | *Paired T-Test between Perception & Memory* | |
|---|---|---|---|---|---|---|---|---|---|
| | | | | t-statistic | p-value | t-statistic | p-value | t-statistic | p-value |
| **OFA** | 13.924 ± 1.978 | 2.631 ± 0.366 | 24 | 6.049 | <0.001 | 1.826 | 0.040 | 5.562 | <0.001 |
| **FFA1** | 15.626 ± 1.727 | 3.575 ± 0.491 | 24 | 7.13 | <0.001 | 3.288 | 0.002 | 7.329 | <0.001 |
| **FFA2** | 12.167 ± 1.227 | 3.651 ± 0.456 | 24 | 8.315 | <0.001 | 3.703 | <0.001 | 7.278 | <0.001 |
| **LOS** | 16.061 ± 1.231 | 2.547 ± 0.344 | 24 | 11.450 | <0.001 | 1.700 | 0.051 | 12.326 | <0.001 |
| **ITG** | 16.876 ± 1.228 | 3.374 ± 0.301 | 24 | 12.142 | <0.001 | 4.689 | <0.001 | 11.504 | <0.001 |
| **MTG** | 10.057 ± 0.941 | 2.486 ± 0.320 | 24 | 8.606 | <0.001 | 1.635 | 0.058 | 8.331 | <0.001 |
| **PPA** | 22.442 ± 1.192 | 10.698 ± 1.048 | 24 | 17.195 | <0.001 | 8.332 | <0.001 | 10.381 | <0.001 |
| **OPA** | 18.656 ± 1.122 | 11.421 ± 1.120 | 24 | 14.881 | <0.001 | 8.446 | <0.001 | 5.205 | <0.001 |

**Title:** Multiple forms of sensory reinstatement in category-selective cortex

**Table S2.** One-Tailed Paired T-Test between Top 10% Preferred Category Uncontrasted Beta Estimates and Each Other Category In Perception and Memory

| *ROI* | *Category Uncontrasted Beta Estimate Mean $\pm$SEM (Preferred Category in Bold)* | | | *DF* | *Perception Paired T-Test* | | *Memory Paired T-Test* | |
|---|---|---|---|---|---|---|---|---|
| | Category | Perception | Memory | | t-statistic | p-value | t-statistic | p-value |
| **OFA** | **Face** | **2.743±0.221** | **0.686±0.124** | 24 | - | - | - | - |
| | Scene | 1.623±0.207 | 0.371±0.074 | | 5.999 | <0.001 | 4.270 | <0.001 |
| | Body | 2.330±0.226 | 0.485±0.104 | | 2.282 | 0.016 | 3.696 | <0.001 |
| | Object | 2.184±0.216 | 0.514±0.110 | | 2.831 | 0.005 | 2.903 | 0.004 |
| **FFA1** | **Face** | **3.170±0.244** | **0.646±0.087** | 24 | - | - | - | - |
| | Scene | 2.531±0.370 | 0.333±0.076 | | 2.236 | 0.017 | 5.281 | <0.001 |
| | Body | 2.682±0.290 | 0.490±0.088 | | 2.675 | 0.007 | 3.839 | <0.001 |
| | Object | 2.537±0.316 | 0.431±0.083 | | 2.677 | 0.007 | 3.164 | 0.002 |
| **FFA2** | **Face** | **1.769±0.159** | **0.387±0.037** | 24 | - | - | - | - |
| | Scene | 0.835±0.106 | 0.094±0.036 | | 6.529 | <0.001 | 6.453 | <0.001 |
| | Body | 1.265±0.136 | 0.212±0.044 | | 4.119 | <0.001 | 4.759 | <0.001 |
| | Object | 0.996±0.131 | 0.170±0.050 | | 5.782 | <0.001 | 4.625 | <0.001 |
| **LOS** | **Body** | **2.967±0.276** | **0.230±0.034** | 24 | - | - | - | - |
| | Scene | 0.960±0.120 | 0.035±0.034 | | 8.330 | <0.001 | 4.025 | <0.001 |
| | Face | 1.711±0.245 | 0.136±0.039 | | 7.388 | <0.001 | 3.370 | 0.001 |
| | Object | 2.336±0.244 | 0.142±0.036 | | 5.742 | <0.001 | 2.715 | 0.006 |
| **ITG** | **Body** | **3.123±0.198** | **0.398±0.056** | 24 | - | - | - | - |
| | Scene | 1.033±0.099 | 0.152±0.053 | | 12.022 | <0.001 | 6.651 | <0.001 |
| | Face | 2.011±0.189 | 0.247±0.059 | | 7.960 | <0.001 | 5.158 | <0.001 |
| | Object | 2.571±0.179 | 0.385±0.072 | | 7.267 | <0.001 | 0.378 | 0.354 |
| **MTG** | **Body** | **1.571±0.169** | **0.246±0.045** | 24 | - | - | - | - |
| | Scene | 0.339±0.086 | 0.135±0.049 | | 8.665 | <0.001 | 2.984 | 0.003 |
| | Face | 0.989±0.129 | 0.115±0.042 | | 6.550 | <0.001 | 2.751 | 0.006 |
| | Object | 1.288±0.158 | 0.166±0.045 | | 3.047 | 0.003 | 3.571 | <0.001 |
| **PPA** | **Scene** | **2.764±0.184** | **0.899±0.116** | 24 | - | - | - | - |
| | Body | 0.272±0.086 | -0.129±0.031 | | 15.131 | <0.001 | 8.423 | <0.001 |
| | Face | -0.214±0.068 | -0.212±0.035 | | 15.035 | <0.001 | 8.687 | <0.001 |
| | Object | 0.945±0.117 | -0.034±0.033 | | 14.198 | <0.001 | 8.565 | <0.001 |
| **OPA** | **Scene** | **1.925±0.125** | **0.806±0.083** | 24 | - | - | - | - |
| | Body | 0.745±0.110 | -0.071±0.056 | | 10.191 | <0.001 | 8.055 | <0.001 |
| | Face | 0.062±0.079 | -0.177±0.053 | | 13.687 | <0.001 | 8.734 | <0.001 |
| | Object | 1.240±0.128 | -0.021±0.044 | | 7.659 | <0.001 | 8.630 | <0.001 |

**Title:** Multiple forms of sensory reinstatement in category-selective cortex

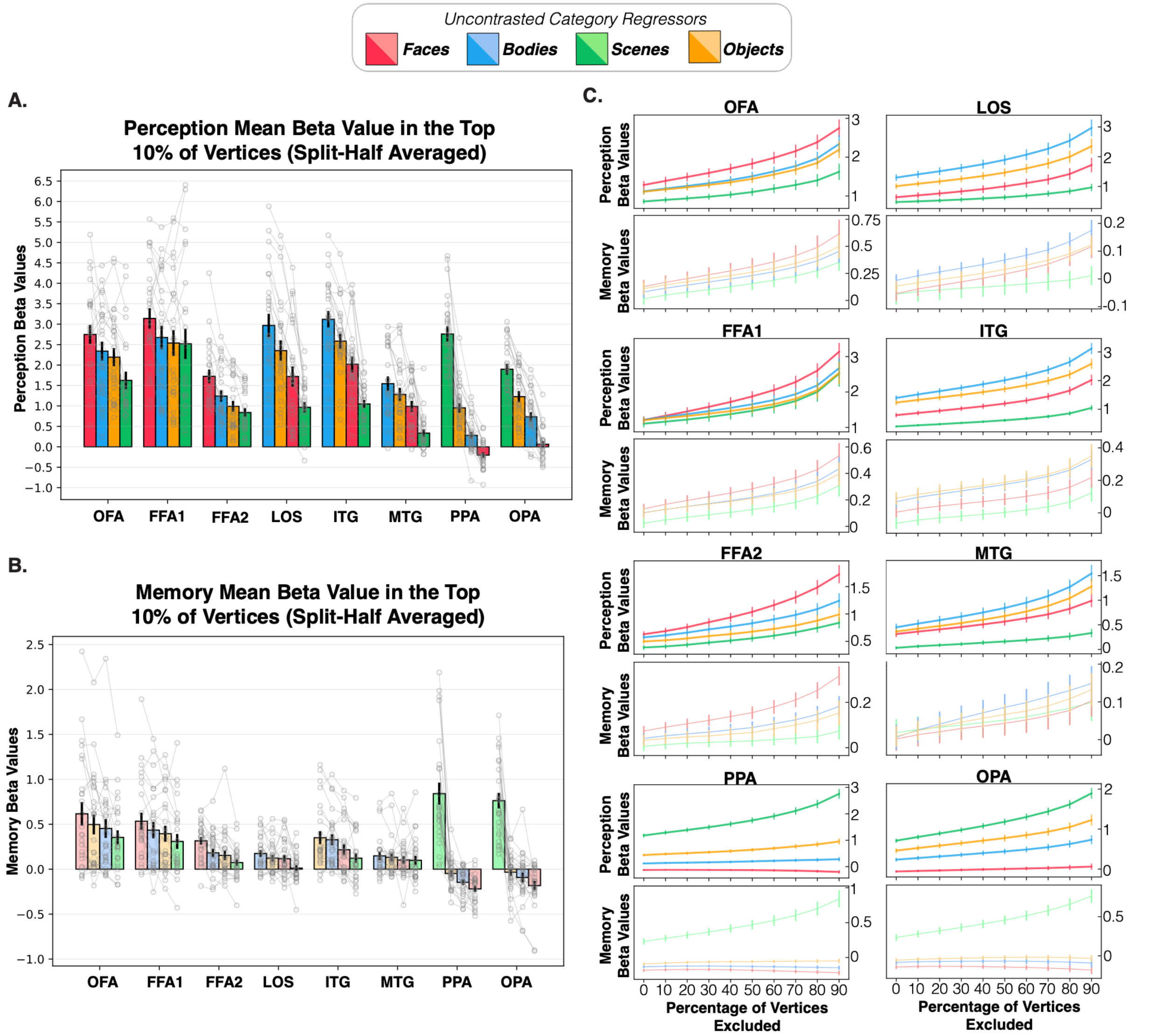


**Figure S1. Category selectivity trends maintained with split-half data**

For each task, the topmost responsive vertices to the preferred category for each ROI were selected using the odd-numbered runs, and the mean beta estimates from those vertices were calculated using the left out even-numbered runs. The converse analysis (select on even-numbered runs, calculate on odd-numbered runs) was also done, with the average between the two plotted here. (A) Mean beta estimates to all four categories in the top 10% most responsive vertices to the preferred category for each ROI during perception, confirming that every ROI responded most strongly to its preferred category. (B) The same analysis during memory, showing a similar trend but weaker overall category selectivity in almost all ROIs (C) Selectivity for the preferred category across decreasing vertex inclusion percentages. Error bars denote SEM.

**Title:** Multiple forms of sensory reinstatement in category-selective cortex

**Table S3.** Difference between Z-Scored Memory Values in Top and Bottom 20% of Perceptually Selective Vertices: One-tailed T-test against Baseline of Zero

| *ROI* | *Number of Participants* | *Mean Difference between Z-Scored Memory Values of Top and Bottom 20% Perception ± SEM* | *DF* | *One-tailed t-test against 0* | |
|---|---|---|---|---|---|
| | | | | t-statistic | p-value |
| **OFA** | 24 | 0.182±0.214 | 23 | 0.848 | 0.203 |
| **FFA1** | 25 | 0.618±0.185 | 24 | 3.334 | 0.001 |
| **FFA2** | 25 | 1.190±0.189 | 24 | 4.946 | <0.001 |
| **LOS** | 23 | 0.393±0.210 | 22 | 1.866 | 0.038 |
| **ITG** | 25 | 0.321±0.168 | 24 | 4.946 | <0.001 |
| **MTG** | 24 | 0.321±0.173 | 23 | 1.847 | 0.039 |
| **PPA** | 25 | 0.907±0.127 | 24 | 7.140 | <0.001 |
| **OPA** | 25 | 0.215±0.183 | 24 | 1.171 | 0.127 |

**Table S4.** Slope of Binned Perception vs. Z-Scored Memory: One-tailed T-test against 0

| *ROI* | *Number of Participants* | *Mean Slope ± SEM* | *DF* | *One-tailed t-test against 0* | |
|---|---|---|---|---|---|
| | | | | t-statistic | p-value |
| **OFA** | 19 | 0.428±0.249 | 18 | 1.716 | 0.052 |
| **FFA1** | 23 | 0.809±0.234 | 22 | 3.454 | 0.001 |
| **FFA2** | 23 | 1.570±0.236 | 22 | 6.642 | <0.001 |
| **LOS** | 23 | 0.554±0.254 | 22 | 2.178 | 0.020 |
| **ITG** | 25 | 1.05±0.208 | 24 | 5.050 | <0.001 |
| **MTG** | 19 | 0.529±0.254 | 18 | 2.080 | 0.026 |
| **PPA** | 25 | 1.037±0.229 | 24 | 6.634 | <0.001 |
| **OPA** | 25 | 0.235±0.156 | 24 | 1.025 | 0.158 |
